# Normalization of Prefrontal Network Dynamics Prevents Cognitive Impairments After Developmental Insult

**DOI:** 10.1101/2025.05.30.657069

**Authors:** Mohamed R. Khalife, Patrick Jasinski, Khalil Abed Rabbo, Mohamed Ouardouz, J. Matthew Mahoney, Rod C. Scott, Amanda E. Hernan

## Abstract

The neurodevelopmental period is highly sensitive; insults during this period impair neural network connectivity, causing lasting cognitive deficits associated with many neuropsychiatric disorders. Medial prefrontal cortex (mPFC) networks subserve flexible behavior, but the mechanisms underlying their disruption after developmental insults remain unclear. We used an early-life seizure (ELS) model to investigate how mPFC networks become impaired and tested whether adrenocorticotropic hormone (ACTH), a clinically relevant neuroprotective peptide, could restore network function. Using *in-vivo* single-unit recordings during baseline and fear extinction learning, we found ELS-induced dysfunction was characterized by reduced neuronal firing, rigid spike-timing, and weakened functional connectivity, all predicting impaired extinction learning. ACTH treatment prevented these deficits, preserving dynamic spike-timing, flexible connectivity, and network organization. Advanced graph neural network modeling identified neuronal features predictive of cognitive outcomes, revealing potential biomarkers broadly relevant to other developmental disorders. These findings highlight fundamental mechanisms of mPFC network dysfunction and emphasize the translational potential of targeting network dynamics to restore cognition in neurodevelopmental disorders.

## Introduction

The prefrontal cortex (PFC) is responsible for information processing that underlies higher order executive function and complex behavior, but our understanding of the neural network underpinnings of such processing is still incomplete. Understanding how the PFC encodes information is crucial because the PFC is particularly vulnerable to disruption that can lead to neurodevelopmental disorders such as autism and ADHD(*1–3*). This is in part because the PFC undergoes prolonged neurodevelopment. The developmental period is a highly vulnerable phase for the brain, during which insults such as maternal immune activation, ischemia, traumatic brain injury, or seizures are associated with enduring disruptions in neural maturation and significant cognitive and behavioral deficits(*4–17*). These insults often manifest as disruptions to the large-scale neural networks underlying specific cognitive functions, highlighting the need for mechanistic understanding and broadly applicable interventions.

Acute recurrent ELS represent one significant model of developmental insult known to disrupt neural network establishment, resulting in persistent cognitive deficits, particularly in learning and memory(*18–20*). We and others have shown that recurrent ELS is associated with impaired cognitive performance across various PFC-dependent tasks(*21–28*). However, quantitative insights regarding large-scale network organization are limited for the mPFC, despite its crucial role in cognitive flexibility and its known vulnerability to developmental insults(*23*, *24*, *29–34*). This critical knowledge gap underscores the importance of interventions capable of broadly preventing or reversing network dysfunctions induced by developmental insults(*26*, *28*, *35*, *36*).

Adrenocorticotropic hormone (ACTH), a melanocortin peptide, is a promising neuroprotective candidate, demonstrating efficacy in mitigating cognitive deficits in PFC- dependent tasks following ELS, independent of seizure parameters(*26*, *37*). Notably, ACTH treatment also demonstrates translational potential due to its clinical accessibility and established safety profile. When a single dose of ACTH is administered during the time of recurrent flurothyl-induced seizures,, it preserves PFC-dependent behaviors into adulthood without altering seizure parameters(*26*, *37*). ACTH influences neural circuits directly via melanocortin-4 receptor (MC4R) expressed in the brain, which is linked to neuroprotection, synaptic plasticity, and astrocyte function(*26*, *37–39*). Yet, the specific network-level mechanisms through which ACTH stabilizes or restores mPFC functional architecture underlying cognitive flexibility remain unexplored.

In this study, we utilize in vivo single-unit recordings in control mice, mice with a history of ELS, and mice with a history of ELS treated with ACTH to prevent behavioral impairments during a fear extinction task that relies on mPFC activity. We use graph-theoretic analyses, and a graph-neural-network (GNN) framework to (1) characterize long-term mPFC network reorganization following recurrent ELS, (2) determine how ACTH acutely administered during developmental insult shifts network evolution trajectories toward normal adult connectivity, and (3) assess for the first time how neuronal features predict individual fear-extinction performance across animals with and without a developmental insult. We find that ELS significantly impairs network cohesion, weakens functional connectivity, reduces clustering coefficients, shifts centrality profiles, and decreases information efficiency. ACTH treatment protects network development, guiding connectivity towards control-like trajectories rather than pathological trajectories induced by developmental insult. Crucially, our GNN model accurately predicts cognitive outcomes from neuronal functional features, offering mechanistic insights into how network dynamics drive fear extinction learning. These results provide a robust network-level mechanistic understanding of cognitive flexibility, demonstrating how network dysfunction emerges after developmental insults and how ACTH intervention alters this trajectory. Importantly, our findings suggest broad applicability to neurodevelopmental disorders characterized by disrupted neural connectivity, thereby expanding therapeutic opportunities across multiple developmental conditions.

## Results

### ACTH Preserves Cognitive Flexibility in Fear Extinction Learning After Early-Life Seizures

The fear conditioning task is a PFC- dependent task(*31*), performed to assess fear learning acquisition and fear extinction. From postnatal day (P) 10- P14, 20 animals underwent 20 flurothyl-induced seizures with a 7% mortality rate. 10 of these animals received vehicle treatment and 10 received ACTH treatment. The 20 seizure animals, in addition to 8 control vehicle-treated animals, performed the task at P55.

On day 1 during fear acquisition, all groups successfully learned the task as shown by increased freezing to the tone (**Fig. 1A**; generalized estimating equations [GEE], main effect of tone p<0.05). There were no significant differences between the groups (GEE, main effect of group p>0.05) and no significant effect of ELS or ACTH treatment (both GEE, p>0.05), indicating similar acquisition performance across groups.

**Figure 1.**
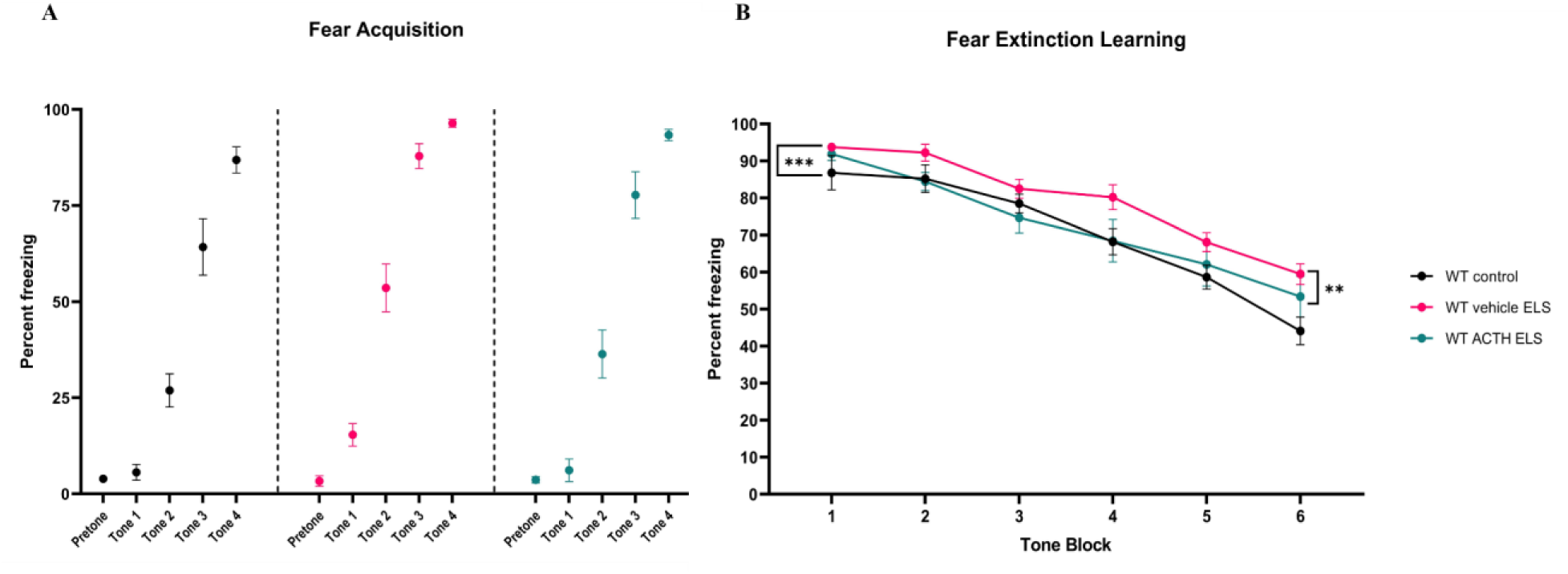
ACTH Improves Fear Extinction Learning Deficit After ELS. All animals across all groups acquired the fear learning paradigm after two tone-shock pairings **(A)**. However, vehicle-treated ELS animals (pink circles) had significant deficits performing the fear extinction part of the task compared to controls (black circles), even after 30 tones without shock pairing. Treatment with ACTH (teal circles) significantly prevented the fear extinction deficit in animals with a history of ELS **(B)**.

On day 2, 24 hours after fear acquisition, the experimental groups performed the fear extinction component of the task. During this task, mice were exposed to a total of 30 pseudo-randomly presented tones without shocks and the percentage of time freezing to the tone throughout the session was assessed. A significant main effect of tone was observed (GEE, p<0.0001), indicating that freezing decreased over time across groups. After adjusting for tone, a significant main effect of group was found (GEE, p = 0.0001), with vehicle-treated ELS mice exhibiting significantly higher freezing than controls (GEE, p = 0.00008), indicating impaired extinction. In contrast, ACTH-treated ELS mice froze significantly less than vehicle-treated ELS mice (GEE, p = 0.009), and were not significantly different from controls (**Fig. 1B**; GEE, p = 0.403), indicating preserved extinction learning.

### ACTH Normalizes Neuronal Firing Rates at Baseline After Early-Life Seizures

8 control, 10 vehicle-treated ELS, and 10 ACTH-treated ELS animals were implanted with single-unit tetrodes and recorded at baseline when they were not engaged in any behavioral task. Single neuron recording shows a significant difference in the firing rate between the control (1.75 Hz, 95% CI [1.1714, 2.3210]) and vehicle-treated ELS group (0.98 Hz, 95% CI [0.86, 1.1042]) (GEE, p=0.01). ACTH normalized the firing rate (1.73 Hz, 95% CI [1.15, 2.32]) compared to the vehicle-ELS group (**Fig. 2A**; GEE, p=0.01). There was no significant difference between the control and ACTH-treated group (GEE, p=1.0). Further, there were no significant differences in the inter-spike intervals between the groups (**Fig. 2B**; control vs ELS vehicle, GEE, p=0.6 and ELS vehicle vs ELS ACTH, GEE, p=0.8 and control vs ELS ACTH, GEE, p=0.9).

**Figure 2.**
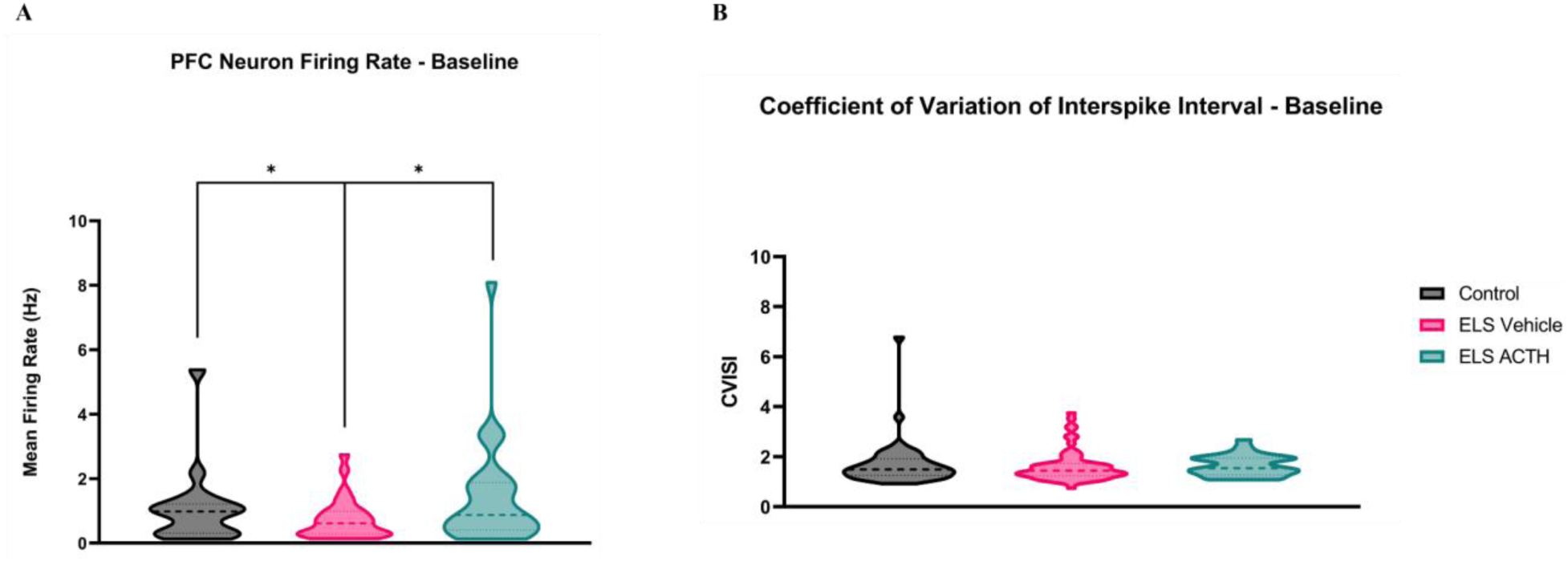
ACTH Normalizes Baseline Firing Rates of Prefrontal Cortex (PFC) Neurons at Baseline. PFC neurons from animals with a history of ELS showed a significant reduction in the mean firing rate compared to control animals. Treatment with ACTH normalized the firing rate deficit observed after ELS **(A)**. The coefficient of variation of interspike intervals did not differ significantly among control, ELS, and ACTH-treated groups **(B)**.

### ACTH Protects Spike-Timing Flexibility in the mPFC After Early-Life Seizures

Neurons rely on the dynamic modulation of their firing probabilities to support complex cognitive processes such as learning and memory(*25*, *40*, *41*). This plasticity in spike timing, often referred to as temporal coding, enables neurons to adapt their firing based on prior spiking activity. To model this temporal coding, we used a generalized linear model (GLM) that incorporates a neuron’s past spiking history and refractory properties to estimate instantaneous firing probability(*42–44*). From the GLM, we captured the properties of fine spike timing using a post-spike filter (PSF) for each neuron, which models the temporal pattern of a neuron’s likelihood of firing after an initial spike. Next, we applied principal component analysis (PCA) on all PSFs to identify the major dimensions of variation in firing patterns, focusing on the first principal component (PC1), which explained the greatest variance across neurons and was then used as a parameter in our statistical analyses. This gives us a robust, statistically principled, and data driven parameter to quantify fine spike timing.

Mean PSFs show group differences in the temporal shape of neuronal firing **(Fig. 3A)**. Vehicle-treated ELS neurons showed two firing peaks at approximately 42 ms and 120 ms after a spike (**Fig. 3A, middle panel**). However, both Control (**Fig. 3A, top panel**) and ACTH-treated ELS (**Fig. 3A, bottom panel**) neurons showed an initial peak around 42 ms and a peak at around 120ms but with decreased amplitude and a smoother decay compared to the ELS vehicle group. To further illustrate these differences, heatmaps of post-spike filters sorted by PC1 score are provided in **Supplementary Figure 1**.

**Figure 3.**
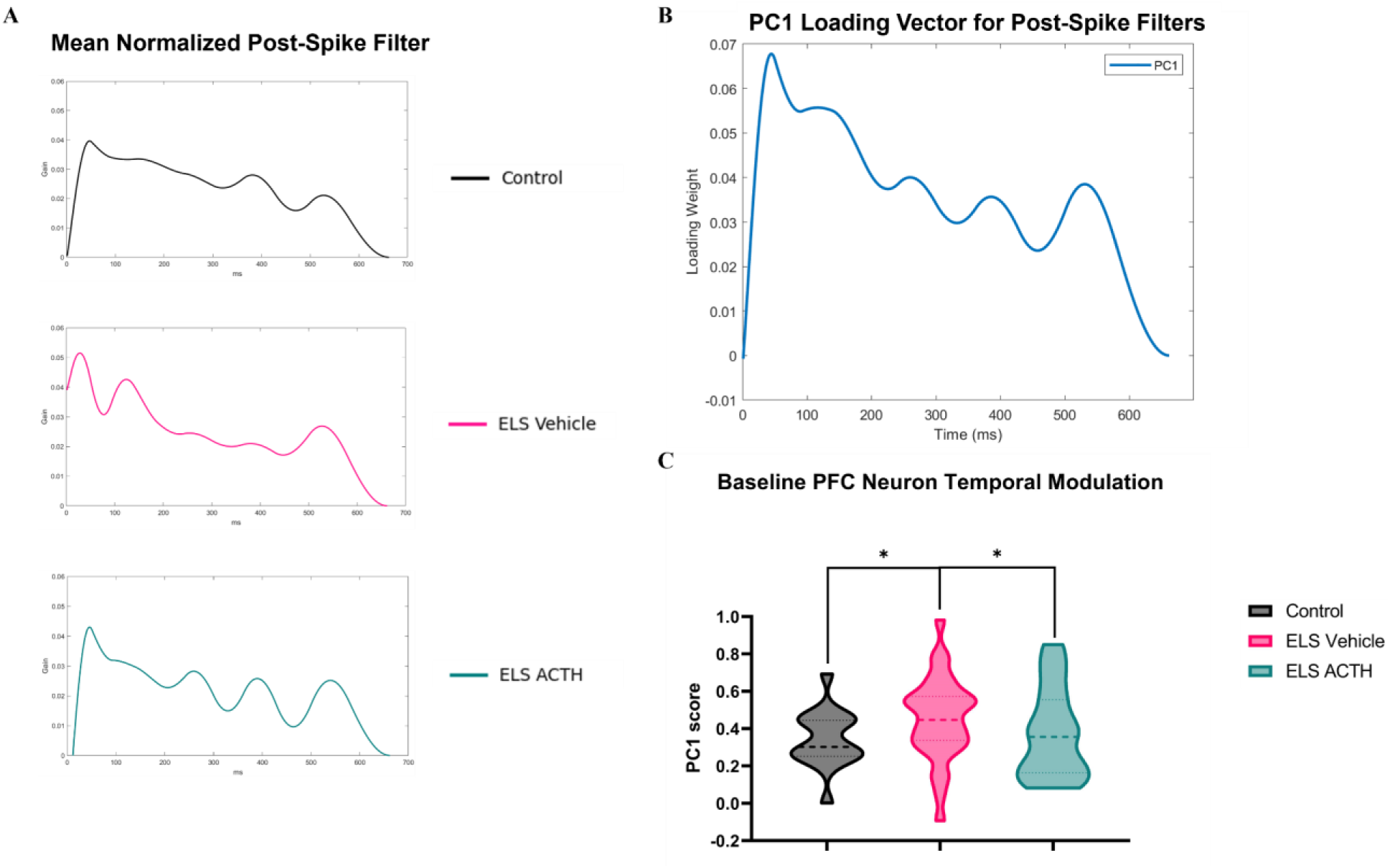
ACTH Normalizes mPFC Neuron Spike Timing Flexibility After ELS. Mean normalized PSF curves illustrate the probability of a neuron refiring over time after an initial spike. Vehicle-treated ELS neurons (pink line) displayed prominent peaks at around 42 ms and 120 ms. However, control (black line) and ACTH-treated (teal line) neurons showed smoother temporal profiles, with a peak at approximately 42 ms that gradually tapers **(A)**. The first principal component (PC1) loading vector from the PCA on all individual PSFs reveals pronounced peaks near 42 and 120 ms **(B)**. PCA analysis on the PSFs reveals that the vehicle-treated ELS group exhibits significantly higher PC1 scores compared to the control group. ACTH treatment showed reduced scores to near-control levels, indicating a preservation of the firing repertoire disrupted by ELS **(C)**.

As expected, the PC1 loading vector captured the same firing variation as seen on the PSF, an early peak at around 42 ms and a delayed rebound at around 120 ms **(Fig. 3B)**. The early peak corresponds to beta-range modulation (around 24 Hz), while the delayed peak aligns with theta-range activity (around 8.3 Hz). Notably, all groups exhibited this temporal structure, but the magnitude and prominence of these features differed significantly across the groups. ELS vehicle neurons exhibited stronger loading on both peaks, particularly the early beta component, indicating hypersynchrony and temporally rigid entrainment to fixed firing windows.

These dynamics were captured by the PC1 score which quantifies the neuron’s alignment with the principal timing pattern seen in the loading vector. Vehicle-treated ELS neurons exhibited significantly higher PC1 scores **(Fig. 3C)** compared to Control (GEE, p = 0.037) and ACTH-treated ELS mice (GEE, p = 0.025), reflecting more rigid action potential timing. In contrast, ACTH treatment preserved flexible firing patterns with PC1 scores at near-control levels (GEE, p = 0.81), indicating preserved firing timing flexibility. These findings show that ACTH mitigates the ELS-induced shift toward a more rigid spike timing pattern, normalizing the temporal coding deficits in the mPFC at baseline.

### Population Dynamics Do Not Differ Between Groups at Baseline

Population dynamics in the PFC are critical for supporting complex cognitive tasks such as decision-making, working memory, and executive control(*45–49*). These tasks rely heavily on population coding where groups of neurons encode stimuli or task-related demands. Disruption of population coding could therefore compromise the neural representation of cognitive demands, potentially underlying the cognitive impairments following ELS. To evaluate if such network disruptions occur under baseline conditions with low cognitive demand, we built graphs from neuronal co-firing and assessed several well-established population-level network metrics. These metrics included edge weight, weighted degree centrality, weighted clustering coefficient, eigenvector centrality, global efficiency, and local efficiency, each quantifying distinct but complementary aspects of neural network function. Edge weights represent functional connectivity between neurons **(Fig. 4A)**. Weighted degree centrality assesses a neuron’s overall connectivity strength within the network **(Fig. 4B)**, whereas the weighted clustering coefficient measures the tendency of neurons to form interconnected cohesive clusters **(Fig. 4C)**. Eigenvector centrality quantifies the influence of a neuron within a network based on the importance of its connections **(Fig. 4D)**. Global efficiency quantifies the overall ability of a network to transfer information **(Fig. 4E)**, and local efficiency measures the robustness of information transfer within local neuron neighborhoods **(Fig. 4F)**.

**Figure 4.**
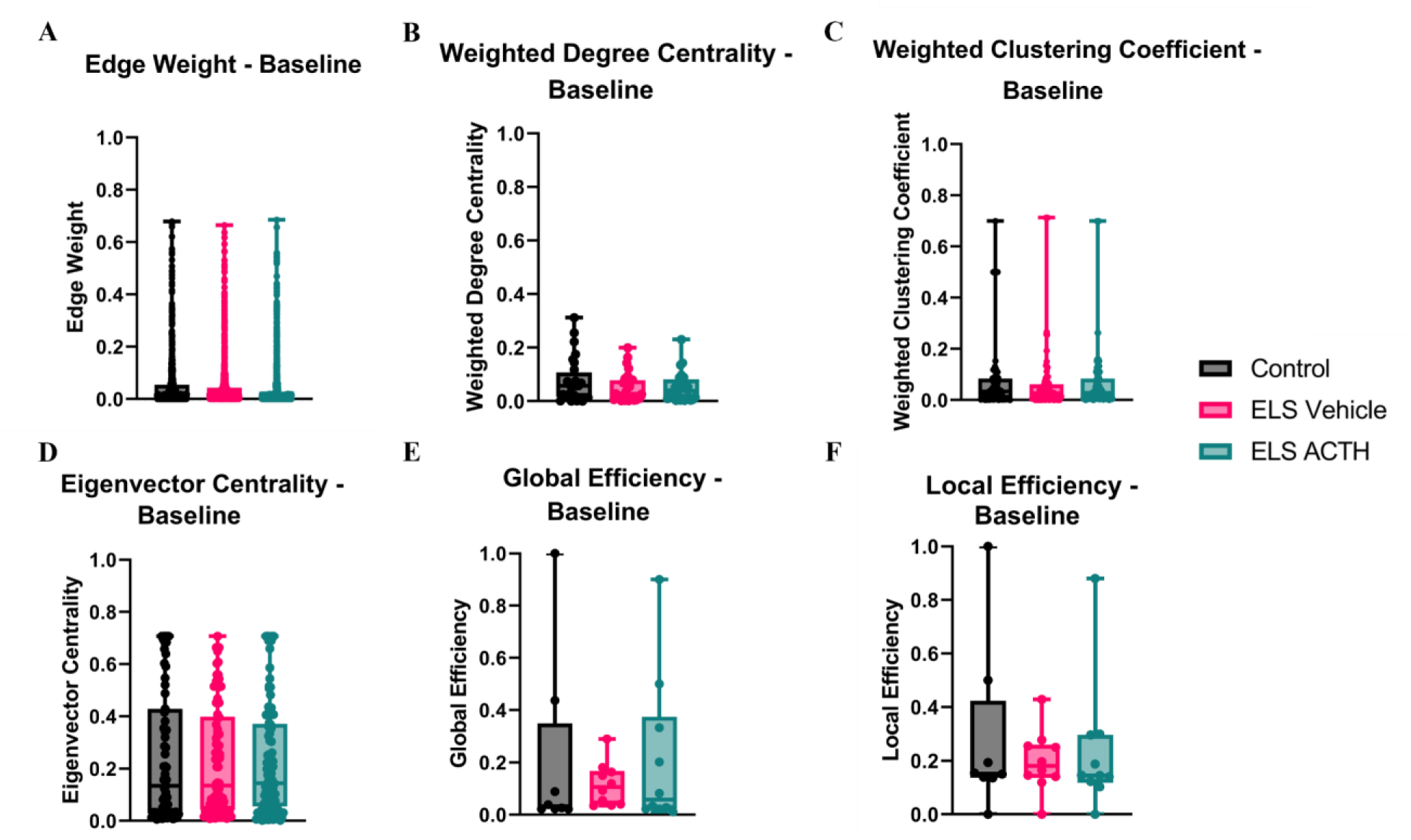
Similar Baseline Connectivity Across Neural Networks From All Experimental Groups. Analysis of PFC neural networks at baseline revealed no significant differences between control, ELS vehicle-treated, and ELS ACTH-treated animals across multiple network metrics. Edge weights **(A)**, weighted degree centrality **(B)**, and weighted clustering coefficient **(C)** showed no significant differences between the groups, suggesting no differences in functional connectivity, node importance, or network clustering propensity, respectively. Similarly, eigenvector centrality **(D)**, global efficiency **(E)**, and local efficiency **(F)** were not significantly different between the groups, indicating no difference in the network integration and communication efficiency at baseline. Collectively, these results suggest that network properties of PFC neurons are similar across all experimental groups at rest.Figure 5**. ACTH Treatment Preserves Neuronal Rate Encoding of Tone During Extinction Learning.** Peri-stimulus time histograms (PSTHs) of firing rate responses to tones during fear extinction learning in Control **(A)**, ELS Vehicle **(B)**, and ELS ACTH **(C)** groups. PSTHs are shown for the first 10 tones (upper panels) and the last 10 tones (lower panels) of the session. Yellow shading indicates tone presentation, and light blue shading indicates the pretone baseline in the first segment. Green bars represent firing during tone periods, and blue bars represent firing during non-tone periods. Quantification of firing rate responses to tone presentations showed significantly impaired tone responsiveness in vehicle-treated ELS neurons, which was restored by ACTH treatment during extinction learning **(D,E)**. Data are presented as mean ± 95% confidence intervals (CIs).

Our analysis revealed no significant differences between the groups in any of these baseline network metrics (GEE, all p > 0.05; **Fig. 4A–F**). The absence of baseline differences indicates that ELS-related population coding abnormalities are not detectable under resting conditions alone.

### ACTH Normalizes Neuronal Response Dynamics During Fear Extinction

We next explored how the brain network changes during fear extinction learning by quantifying the evolution of firing and co-firing dynamics throughout the fear extinction session. To examine firing rate dynamics in response to tones, we quantified the neuron-tone response, defined as the change in neuronal firing rate during tone presentation relative to the pre-tone baseline. Peri-stimulus histograms (**Fig. 5A-C**) were used to visualize these firing rate changes in response to tones, focusing on the first and last 10 tones of the 30-tone session (highlighted in yellow) compared to the pre-tone baseline (light blue). During the initial phase of extinction, the vehicle-treated ELS group exhibited a significant reduction in the neuron-tone response compared to the control group (87.97% vs. −6.52%, 95% CI [−2.12, 178.07] vs. [−13.00, −0.05]; GEE, p = 0.04). However, ACTH treatment significantly increased the response compared to the vehicle-treated ELS group (15.53% vs. −6.52%, 95% CI [6.26, 24.82]; GEE, p = 0.0001).

**Figure 5.**
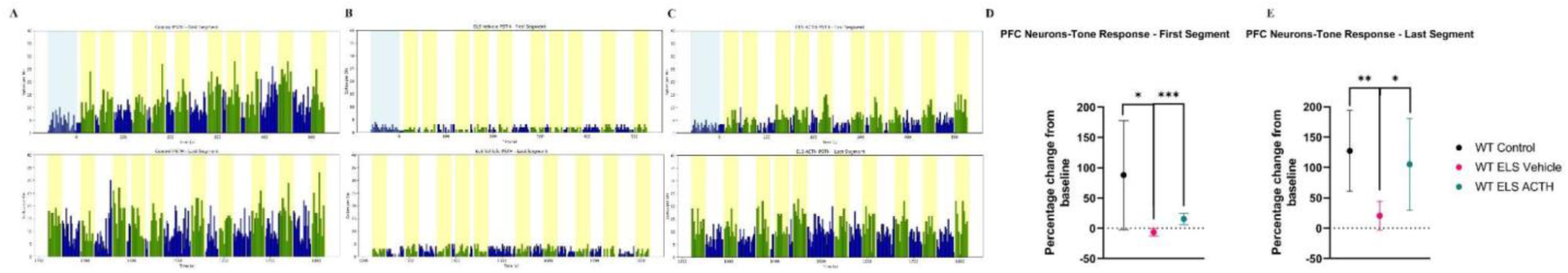
ACTH Treatment Preserves Neuronal Rate Encoding of Tone During Extinction Learning. Peri-stimulus time histograms (PSTHs) of firing rate responses to tones during fear extinction learning in Control **(A)**, ELS Vehicle **(B)**, and ELS ACTH **(C)** groups. PSTHs are shown for the first 10 tones (upper panels) and the last 10 tones (lower panels) of the session. Yellow shading indicates tone presentation, and light blue shading indicates the pretone baseline in the first segment. Green bars represent firing during tone periods, and blue bars represent firing during non-tone periods. Quantification of firing rate responses to tone presentations showed significantly impaired tone responsiveness in vehicle-treated ELS neurons, which was restored by ACTH treatment during extinction learning **(D,E)**. Data are presented as mean ± 95% confidence intervals (CIs).

By the end of the session, firing rate dynamics had evolved across groups (**Fig. 5A-C, lower panels**). However, a persistent deficit in the vehicle-treated ELS group was evident by the significantly reduced responses compared to the control group (127.44% vs. 20.61%, 95% CI [60.99, 193.91] vs. [−3.06, 44.29]; GEE, p = 0.003). ACTH treatment normalized these firing rate responses compared to the vehicle-treated group (105.32% vs. 20.61%, 95% CI [29.60, 181.04]; GEE, p = 0.04), indicating that ACTH normalizes the firing rate encoding of the tone observed following ELS. These results indicate that while tone-responsive neuronal firing rate increased throughout extinction in control animals that learned, neurons from the vehicle-treated ELS group did not change their firing rate over the course of the task, whereas ACTH treatment normalized firing rate responses.

### ACTH Maintains Network Adaptability During Learning

Investigating population dynamics is pivotal for elucidating neural mechanisms underlying cognitive tasks(*45–49*). We investigated the same population metrics used at baseline at the beginning and end of fear extinction.

At the beginning of the fear extinction task, most network metrics, including edge weight, clustering coefficient, eigenvector centrality, and local and global efficiency, showed no significant differences across groups (GEE, p>0.05; **Fig. 6A, C-F**), corroborating our findings at baseline and suggesting initially comparable network structures before learning begins. However, weighted degree centrality was significantly lower in vehicle-treated ELS mice compared to control and ACTH-treated mice **(Fig 6B)**, indicating early deficits in overall connectivity strength. During the last segment of the task, the PFC networks in controls and ACTH-treated animals had strengthened over baseline, underscoring the learning that had occurred during the session. However, neural networks from vehicle-treated ELS mice **(Fig. 6A-F)** did not evolve over the course of the session, characterized by lower functional connectivity, reduced local cohesiveness, compromised influential node connectivity, and impaired global and local network efficiency. The preservation of these metrics by ACTH highlights its ability to protect network-level function critical for cognitive flexibility. Results summarized in **Supplementary Table 1.**

**Figure 6.**
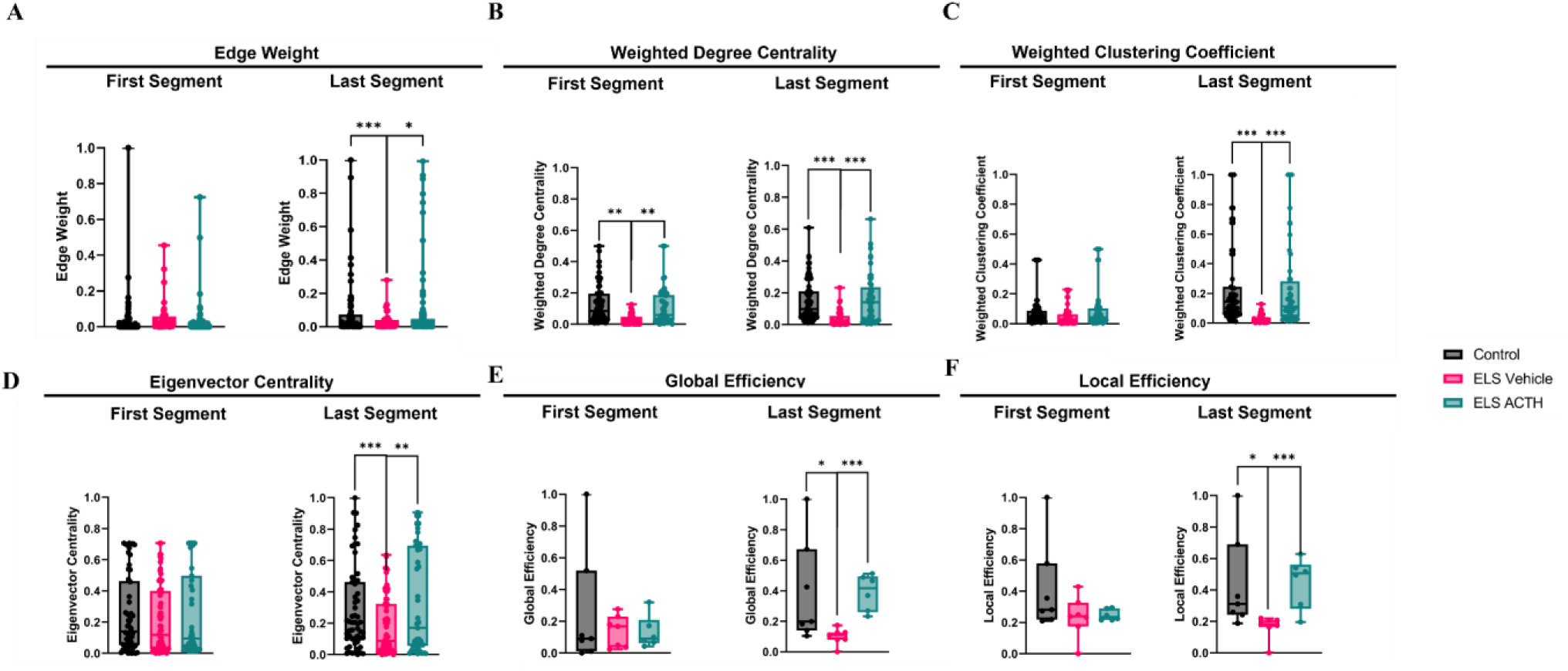
ACTH Normalized Network Connectivity During Learning, Enabling Effective Local And Global Network Communication Essential for Adaptive Behavior. Edge Weight represents the strength of functional connectivity between PFC neurons. No significant group differences (control in black, vehicle-treated ELS in pink, ACTH-treated ELS in teal) were observed in the first segment of the extinction session, but by the last segment, edge weights in the control and ACTH-treated ELS groups were significantly higher than in the vehicle-treated ELS group **(A)**. Weighted Degree Centrality assesses total connectivity, accounting for both the number and strength of a neuron’s connections. Vehicle-treated ELS mice showed lower degree centrality relative to controls and ACTH-treated ELS mice in both the first and last segments, indicating persistent connectivity deficits in the vehicle-treated group that were rescued by ACTH **(B)**. Weighted Clustering Coefficient measures local network cohesiveness. Although no differences emerged in the first segment, the control and ACTH-treated ELS groups exhibited significantly higher clustering coefficients than vehicle-treated ELS mice by the end of the session, suggesting enhanced local connectivity with ACTH treatment **(C)**. Eigenvector Centrality quantifies each neuron’s influence by weighting its connections to other highly connected neurons. All groups showed similar centrality in the first segment, but the control and ACTH-treated ELS groups exhibited a significant increase by the last segment compared to vehicle-treated ELS animals, suggesting that ACTH normalizes centrality properties critical for adaptive network function **(D)**. Global Efficiency reflects the ease of information flow throughout the entire network. In the first segment, no group differences were detected, however global efficiency in the vehicle-treated ELS group was significantly lower than that of control and ACTH-treated ELS mice by the end of the session, underscoring broader integration deficits in the vehicle-treated ELS group **(E)**. Local Efficiency is a measure of the network’s fault tolerance and communication within local neighborhoods of nodes. Similar local efficiency was observed initially across all groups, but the vehicle-treated ELS group’s local efficiency was significantly lower than that of controls and ACTH-treated ELS mice in the last segment **(F)**, suggesting that ACTH treatment enables effective local communication and resilience in PFC circuits.

To further explore relationships among these network metrics, correlation analyses were conducted separately for the initial and final segments of fear extinction **(Supplementary Figure 2)**. During the first segment, network metrics showed modest correlations. However, correlations among network metrics became more pronounced in the last segment, with strong positive correlations emerging between mean edge weight and eigenvector centrality (r=0.93). Eigenvector centrality was strongly correlated with global efficiency (r=0.77). Additionally, global and local efficiencies were highly correlated (r=0.84). These enhanced correlations suggest that as extinction progressed, distinct network properties became increasingly coupled or coordinated, reflecting integrated changes from single-neuron interactions like edge weights to higher-order network properties like efficiency and centrality. This strengthened coupling among metrics likely represents an adaptive reconfiguration of network communication as the result of neuronal plasticity, facilitating efficient neuronal processing essential for successful extinction learning.

### Evolution of Network Dynamics During Baseline and Learning

Baseline network analysis revealed no significant differences in PFC network properties between control, ELS vehicle-treated, and ELS ACTH-treated groups. Baseline radar plot shows no significant group-level differences in terms of connectivity, clustering, integration, or efficiency, suggesting that network properties are initially similar when animals are at rest (**Fig. 7A**).

**Figure 7.**
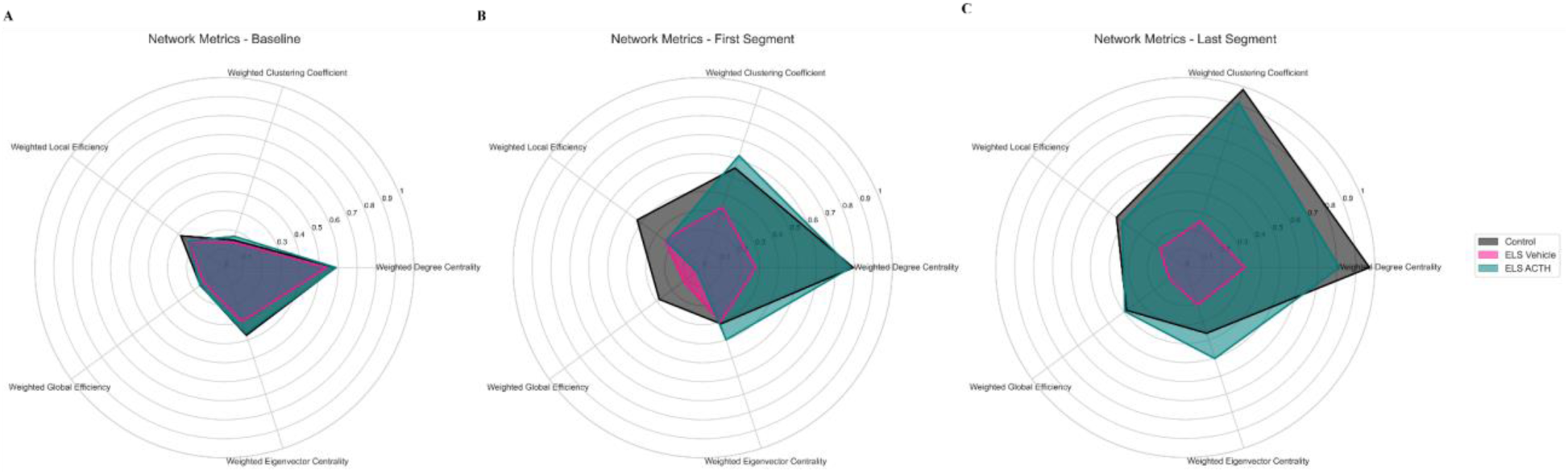
Network Properties Evolved Significantly in Control and ACTH-Treated Groups Across Extinction, In-Contrast to Persistent Deficits Observed in Vehicle-Treated ELS Networks. Radar plots depicting normalized network metrics for control (black), ELS vehicle-treated (pink), and ELS ACTH-treated (teal) groups during baseline **(A)**, the early learning phase of fear extinction (first segment, **B**), and the late learning phase (last segment, **C**). Metrics include edge weight, weighted degree centrality, weighted clustering coefficient, eigenvector centrality, global efficiency, and local efficiency. At baseline, no significant differences were observed between groups, indicating similar network properties during rest **(A)**. In the first segment, significant reductions in connectivity-related metrics were observed in the vehicle-treated ELS group compared to the control and ACTH-treated groups **(B)**. In the last segment, control and ACTH-treated ELS groups exhibited enhanced network integration and connectivity, compared to the vehicle-treated ELS group **(C)**.

During the fear extinction task, the network metrics indicate that at the beginning of the session before any learning has occurred, network properties are comparable across groups, except for significant reductions in overall connectivity in the vehicle-treated ELS group (**Fig. 7B**). As the fear extinction task progressed and the animals learned, differences in network metrics emerged.

Radar plot analysis indicated enhanced network connectivity, integration, cohesiveness, and efficiency in control and ACTH-treated groups compared to persistent deficits in vehicle-treated ELS mice (**Fig. 7C**), suggestive of diminished neural flexibility and adaptation during fear extinction underlying their learning deficit.

### Prediction of Behavioral Outcomes from Neuronal Features in a Graph Neural Network Model

To move beyond traditional correlation-based analyses and assess how neural dynamics actively shape behavior, we implemented a Graph Neural Network (GNN). We then trained an optimized Graph Attention Network (GAT), which incorporates both node-level neuronal features and dynamic connectivity patterns, to predict freezing behavior. This model not only captures the rich structure of population dynamics, but also offers a powerful tool to test whether and how features contribute to behavior. In doing so, it allows us to move closer to understanding network activity as a mechanistic driver of cognition, rather than a correlation.

The optimized GAT model effectively predicted freezing behavior during both first and last segments of fear extinction. The GAT achieved higher predictive performance compared to the linear regression baseline; during the first segment, the GAT model achieved an R² of 0.54 **(Fig. 8A)**, outperforming the linear regression’s R² of −0.24 **(Fig. 8C)**. In the last segment, the GAT showed further enhanced predictive capability, reaching an R² of 0.81 **(Fig. 8B)** compared to 0.28 by linear regression **(Fig. 8D)**. Analysis of model feature importance using GNNExplainer identified maximum ISI, CV of ISI, and firing rate as key predictors during initial extinction, aligning with GEE findings. During the last segment, firing rate, maximum ISI, and PC scores emerged as critical predictors, also consistent with GEE findings. In contrast, linear regression identified fewer and weaker associations, underscoring the GAT’s superior ability to capture biologically relevant neuronal dynamics.

**Figure 8.**
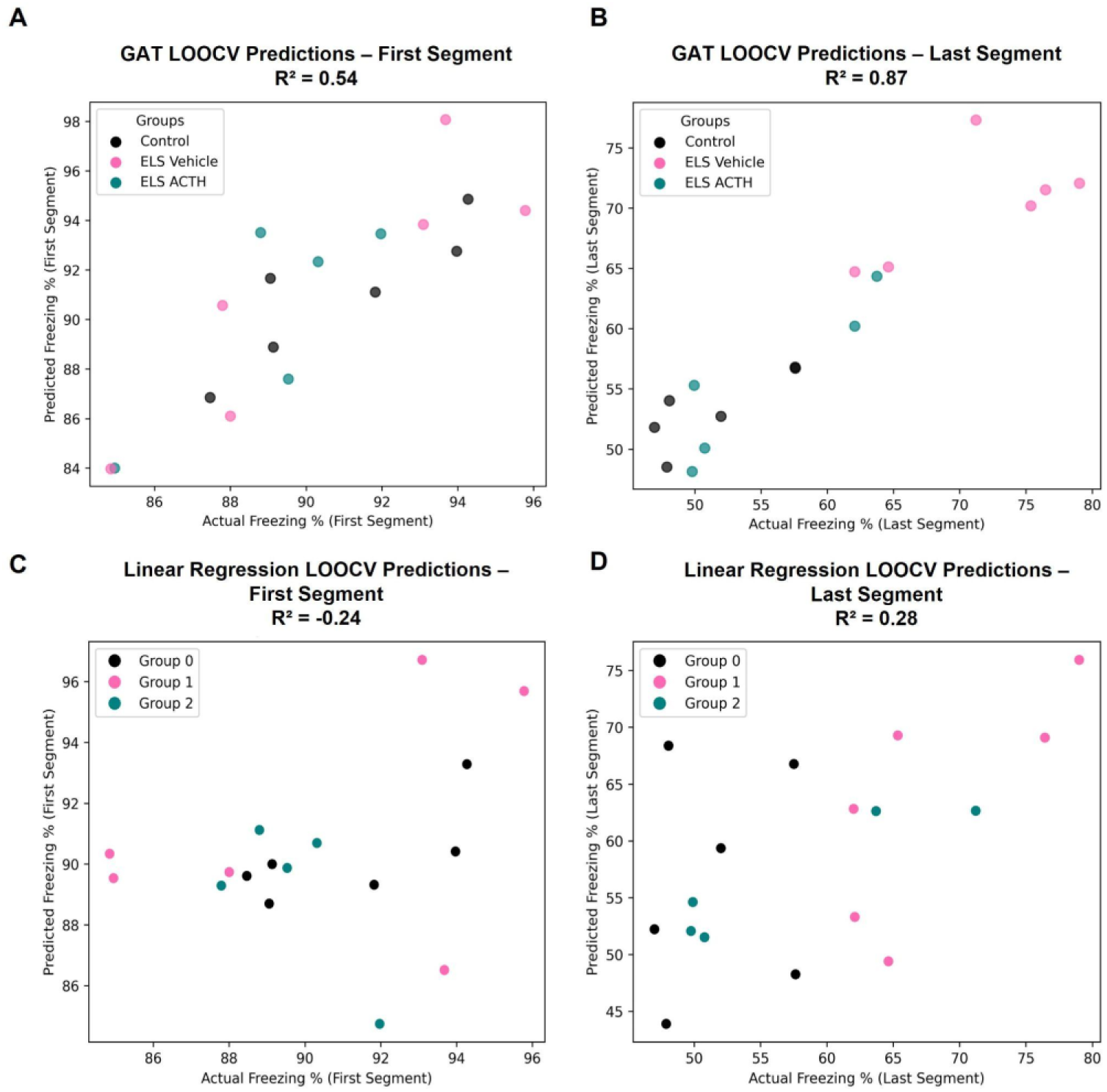
Optimized Graph Attention Network Significantly Outperformed Linear Regression, Demonstrating Superior Predictive Power And Identifying Critical Neuronal Features Contributing to Extinction Behavior. Scatter plots show predicted versus actual freezing percentages for the first **(A&C)** and last **(B&D)** segments of the fear extinction task. Predictions were generated using Leave-One-Out Cross-Validation (LOOCV). The optimized Graph Attention Network model outperformed linear regression by capturing node-level neuronal features and network connectivity, achieving an R² of 0.54 during the first segment and 0.87 during the last segment **(A-B)**. Linear regression predictions were notably less accurate, resulting in a negative R² (−0.24) during the first segment and an R² of 0.28 during the last segment **(C-D)**. Colors represent experimental groups: Control (black), Vehicle-treated ELS (pink), and ACTH-treated ELS (teal).

These findings demonstrate that the optimized GAT model effectively bridges the analytical gap between the linear regression approach, predictive but lacking node-level resolution, and the GEE method, node-level analytical power without predictive capacity, offering robust predictions alongside interpretable insights into network-level neural mechanisms underpinning fear extinction behavior.

## Discussion

We set out to characterize the neural network underpinnings of fear extinction learning using a disease model with impaired learning, and animals that were treated with ACTH to prevent impairments. Early-life seizures (ELS), occurring during critical developmental windows, exert lasting effects on cortical neural networks leading to persistent cognitive deficits characterized by impaired learning, memory, and executive functions(*50–53*). We have previously shown that recurrent flurothyl-induced ELS produces lasting alterations in short-term plasticity within mPFC microcircuits, most prominently in layers II/III to V and layer V to V networks, pointing to a disruption of synaptic plasticity after ELS(*24*). In parallel, our earlier work demonstrated that ACTH’s neuroprotective effect is dependent on MC4R signaling(*37*), and that ACTH but not a corticosteroid selectively normalizes gene-expression pathways related to synaptic plasticity and metabolic homeostasis following ELS(*54*). Notably, MC4R activation through ACTH normalizes astrocyte protein expression and astrocytic activation after ELS(*37*). However, behavioral outcomes depend on the integrity of network-level functional connectivity; how ELS altered functional connectivity underlying behavioral impairment and whether ACTH normalized the network or led to some other compensatory change to subserve normal learning remains unknown. In this study, we show that a single dose of ACTH administered before the first seizure each day over five-days of flurothyl-induced seizures (5 doses of ACTH total), shifts the trajectory of mPFC network evolution towards control-like adult topology, preserving functional connectivity at both the single-neuron and population levels during baseline and fear-extinction learning, thus protecting cognitive flexibility against ELS-induced deficits. Critically, this is despite seizures occurring at the same rate in all groups.

We found significant alterations in single-neuron firing patterns, temporal fine spike-timing, and network-level connectivity after ELS, each of which limit the cognitive flexibility required for adaptive behaviors such as fear extinction learning. Importantly, ACTH administration effectively prevented these alterations, maintaining firing dynamics and connectivity throughout learning. These findings underscore our central premise, that protecting neural network dynamics during the insult is key to preventing the chronic cognitive impairment that follows early life insults. Our results therefore provide a novel mechanistic framework for understanding how early interventions may broadly prevent cognitive deficits stemming from diverse developmental challenges.

Fear extinction learning requires dynamic modulation of neuronal activity within mPFC-basolateral amygdala (BLA) activity to facilitate adaptive suppression of fear responses in the absence of aversive stimuli(*1*, *31*, *55–57*). Dysfunction within these circuits is implicated in pathological conditions such as PTSD, anxiety disorders, and epilepsy, which are characterized by diminished neural flexibility, impaired extinction and learning(*58–63*). Our data suggest that the reduced firing rates and restricted firing repertoire at baseline observed in ELS animals represent critical barriers to the flexible neuronal modulation (i.e. appropriate neural plasticity) necessary for extinction learning. Notably, ACTH treatment preserved intrinsic firing properties, including normalized post-spike filter and PC scores, thus expanding the firing repertoire and facilitating adaptive neuronal responses that are important for extinction learning(*30*, *34*, *44*).

At the single-neuron level, vehicle-treated ELS mice exhibited significantly reduced baseline firing rates and abnormal temporal firing dynamics. These observations align with prior evidence highlighting disrupted intrinsic neuronal dynamics and plasticity following early life seizures(*23*, *24*, *44*). Notably, temporal coding, viewed as the variability and precision of action potential timing, supports cognitive processes such as memory, attention, and behavioral flexibility(*32*, *45*, *48*, *64–66*). Despite the reduced firing rate, the neurons from vehicle-treated ELS animals display a level of beta and theta modulated firing with two firing peaks at approximately 42 and a more prominent 120 ms rebound peak compared to controls and ACTH-treated ELS animals. Principal component analysis confirmed this pattern, with PC1 loadings capturing both peaks and vehicle-treated ELS neurons displaying significantly higher PC1 scores than the other groups.

Thus, vehicle-treated ELS neurons retained a rigid and stereotyped temporal firing structure, firing at a lower rate but with a fixed refiring window, potentially indicative of a maladaptive form of temporal coordination in neuronal firing at the single neuron level. The early 42 ms peak falls within the beta frequency range (20–30 Hz), which has been implicated in maintaining current cognitive states, internal representation, and working memory stability(*67–69*). While beta is a normal feature of the prefrontal coding and was present in all groups, the increased expression of this 42 ms peak in the vehicle-treated ELS group may reflect a pathological persistence indicating a failure to update or flexibly shift active states(*67*). Such temporal rigidity is seen in Parkinson’s disease, where excessive beta presence impairs behavioral flexibility and cognitive control(*69*, *70*). Likewise, in healthy humans, elevated beta bursts suppress adaptive learning by reducing behavioral variability in task prediction and overreliance on older memory representations for reward prediction(*71*). The second 120ms peak aligns with theta rhythmicity (4-8 Hz) which is typically associated with cognitive engagement, however the emergence of a theta-modulated refiring at baseline in the vehicle-treated ELS group likely reflects a constrained, inflexible temporal pattern rather than an increased rhythmic firing underlying learning over time.

These findings illustrate that while beta and theta typically support cognitive stabilization and engagement, their exaggerated presence at baseline reflects pathological rigidity rather than adaptive function by locking the system into a persistent state and limiting dynamic cognitive engagement. Prominent presence of both beta and theta convey a maladaptive form of temporal coordination which means that action potentials are locked into narrow time windows, which reduces the neuron’s ability to adapt to changing inputs or participate in task-driven network dynamics. Such rigidity in spike-timing patterns can impair neuronal coding flexibility, limiting adaptive responses to changing environmental demands(*44*, *72–74*). This impairment in coding flexibility has been shown to impact cognitive functions, as flexible spike timing is critical for efficient sensory coding(*74*, *75*), plasticity and learning(*76*), reliable cortical information processing (*64*), motor output precision(*73*), and working memory function(*72*). Hence, the loss of spike-timing flexibility observed after ELS is one of the metrics that predicts the cognitive deficits observed in our model. Early treatment with ACTH preserved both firing rate and spike-timing variability, supporting more variable spike-timing patterns which lower PC1 scores. This supports a more dynamic and flexible neuronal firing repertoire, contributing to improved cognitive performance and facilitating extinction learning.

Interestingly, we saw no group differences in the coefficient of variation of inter-spike interval (CV ISI) during baseline. CV ISI values across all groups were approximately 1, consistent with Poisson-like spike trains, suggesting that neurons fired randomly overall, and that CV ISI fails to detect structured temporal patterns that reflect maladaptive network dynamics. This highlights the added value of GLM-derived PSFs that can reveal hidden temporal constraints that underlie the loss of spike-timing flexibility after ELS. Notably, despite the differences in single-neuron firing properties at baseline, network-level graph metrics such as centrality, local clustering, and efficiency did not differ significantly between the groups. This suggests that cognitive impairments may not emerge from resting network organization alone, but instead from an inability to dynamically reorganize networks in response to task demands.

Since our goal was to characterize firing dynamics during learning, we employed a fear acquisition and extinction paradigm known to rely on mPFC. Behaviorally, all groups acquired the tone-shock association normally during the acquisition phase of the fear conditioning task, confirming intact initial fear learning. However, during extinction, vehicle-treated ELS mice displayed persistently elevated freezing, indicating impaired learning suppression of the conditioned response. In contrast, ACTH-treated ELS mice exhibited significantly reduced freezing throughout the extinction session, suggesting that ACTH protected PFC dynamics necessary for successful fear extinction. This highlights ACTH’s capacity to preserve cognitive flexibility required for adaptive learning following early developmental insults.

Single neuron analysis during extinction revealed that vehicle-treated ELS neurons exhibited impaired responsiveness to task-related stimuli during extinction learning. These neurons showed significantly reduced firing rate increases in response to tone presentations compared to both control and ACTH-treated ELS groups, as seen in the peri-stimulus time histograms. This deficit in tone-evoked response was observed both at the beginning and end of extinction, suggesting a persistent failure of stimulus engagement. In contrast, neurons from ACTH-treated animals showed restored tone-responsiveness, supporting the role of ACTH in preserving both intrinsic and task-evoked activity. These data suggest that temporal rigidity and low baseline activity in ELS neurons are accompanied by a failure to dynamically engage with task events, which further reinforces the idea that these neurons are functionally disconnected from adaptive network processes. Taken together, this suggests that the impairments in network plasticity and adaptability, rather than baseline abnormalities, underlie the observed cognitive deficits. Our findings thus reveal key insights into the functional consequences of disrupted network plasticity, highlighting the importance of maintaining adaptive network dynamics for cognitive resilience.

To understand population-level effects during the fear extinction task, we quantified multiple graph-based metrics including edge weights, weighted degree centrality, clustering coefficients, eigenvector centrality, and global/local efficiencies. These metrics collectively provide detailed insights into different aspects of functional connectivity, network integration, cohesiveness, and communication efficiency(*77*, *78*). Vehicle-treated ELS mice exhibited significant impairments across these metrics during fear extinction, notably showing reduced integration, lower centrality, disrupted local clustering, and diminished global efficiency, which are features consistent with fragmented and inefficient network communication. In contrast, networks built from neurons recorded in ACTH-treated ELS mice exhibited properties similar to control animals, indicating that maintaining these critical network features is important for adaptive fear learning. The protection of local clustering, integration, efficiency and centrality metrics in ACTH-treated animals support enhanced local and global network interactions, which are crucial for effective cognitive performance(*79–81*). These network-level impairments likely emerge from a combination of single-neuron temporal constraints, reduced firing, impaired stimulus responsiveness leading to changes in network plasticity and ultimately poor learning in the task. Together, these results highlight that reduced firing and rigid spike-timing in ELS animals restrict the neuron’s ability to engage dynamically with evolving task demands, contributing to the network-level fragmentation observed during extinction. ACTH treatment prevented these constraints, restoring responsiveness and facilitating neuronal integration, indicating that stimulus responsiveness, firing rate, and spike-timing are interconnected aspects of network adaptability that converge to support flexible cognition. This emphasizes the broad potential for early interventions targeting network plasticity and connectivity to mitigate cognitive deficits across neurodevelopmental disorders.

Next, we asked how the whole landscape of population dynamics relates to extinction learning. We performed correlation analyses among these metrics separately at the initial and final segments of extinction and revealed how each metric captures distinct yet interrelated aspects of network function. At the beginning of extinction, while the behavioral performance of the animals is the same, network metrics displayed moderate to weak correlations, suggesting independence in how different network properties reflect neuronal dynamics during initial task engagement. By the end of extinction, correlations among metrics became stronger, reflecting tighter coordination among network properties. These stronger correlations during the last segment of extinction indicate increased coupling among network properties, reflecting coordinated adaptations across multiple levels, from local neuronal interactions like edge weights to broader network properties like efficiency and centrality. However, despite the global increase in correlations, the radar plots reveal that the vehicle-treated ELS group failed to evolve in the magnitude of these network features during the extinction task. While control and ACTH-treated groups showed increases in network metrics by the end of extinction, the vehicle-treated ELS group remained significantly lower across all of them, highlighting that both network integration and magnitude are required for successful fear extinction. This suggests that successful fear extinction learning relies on coordinated changes across multiple aspects of network organization within the mPFC such as local clustering, connectivity strength, and global efficiency rather than independent single parameter changes. Thus, ACTH’s preservation of these integrated metrics likely facilitates the cohesive neuronal interactions and dynamics reorganization essential for learning(*79–81*). These population-level findings have significant implications, suggesting that early interventions aimed at preserving network integrity could help mitigate long-term cognitive deficits after an early-life insult. Importantly, our approach provides a framework for systematically investigating how interventions influence integrated network properties underlying cognitive resilience.

Finally, to further characterize neuronal contributions and directly link neural dynamics to behavior, we implemented a Graph Neural Network (GNN) model utilizing a Graph Attention Network (GAT) architecture. Unlike traditional statistical approaches such as linear regression, which captures population-level data, or generalized estimating equations (GEE), which provide node-level resolution without predictive capability, our optimized GAT model integrates node-level features and their connectivity to robustly predict behavioral outcomes. Rather than identifying correlations, this model helps us ask a more important question regarding which features of neural activity are actually influencing and predicting behavior, allowing us to bridge the gap between behavior and mechanism to show whether neural dynamics *determine* cognitive outcomes. The GAT model significantly outperformed linear regression in both segments of the task suggesting that the higher R² reflects not just improved model fit but the emergence of biologically structured dynamics over the course of learning, highlighting the value of incorporating complex neuronal interactions into behavior prediction. Feature importance analysis revealed that firing rate, ISI properties, and PSF principal components were strong predictors of freezing behavior during fear extinction, reinforcing that both firing and temporal coding are behaviorally relevant. This demonstrates the utility of the GNN approach with its capacity to integrate neuron-level analytical depth, like GEE, but with predictive power. Hence, this innovative approach not only enhances our mechanistic understanding of network dynamics but also holds promises for future applications such as predicting treatment responses and exploring network dysfunctions in other neurodevelopmental conditions. This methodological advancement thus provides an important translational tool for identifying predictive network biomarkers and developing targeted interventions across various developmental disorders.

Although our study reveals multi-scale mechanisms linking early-life insult to adult cognitive flexibility, it is important to point out several limitations. First, recordings were limited to the mPFC and fear extinction where contributions of interconnected regions like basolateral amygdala and hippocampus remain untested. Second, we modeled only flurothyl-induced seizures, hence generalizability to other developmental insults and to post-insult therapeutic windows requires evaluation. Third, translating our network biomarkers to non-invasive modalities like EEG and fMRI will be critical for clinical application.

In conclusion, our findings demonstrate that ELS produces long-lasting impairments in both spike-timing structure and network connectivity that limit cognitive flexibility during fear extinction learning. Early treatment with ACTH leads to long-lasting preservation of both firing variability and network adaptability, reinforcing the notion that interventions aimed at protecting network-level dynamics can effectively mitigate the long-term cognitive consequences of early developmental insults. By normalizing firing rates, temporal coding, and key connectivity metrics, ACTH facilitates dynamic PFC reorganization essential for adaptive learning. These results reveal how single-neuron alterations propagate to large-scale dysfunction and underscore the therapeutic potential of targeting neural dynamics. The translational potential extends broadly to other neurodevelopmental characterized by disrupted neural connectivity, supporting further exploration of ACTH or other network-focused interventions to mitigate diverse forms of cognitive impairment. Future research should probe MC4R-downstream pathways, test network-level interventions across other neurodevelopmental disorders, and validate the predictive utility of network models. Further application of our graph-based methodologies and GNN predictive modeling will provide valuable insights into the generalizability of network-level biomarkers, enabling more effective interventions for developmental cognitive disorders.

## Materials and Methods

### Animals

Male and female C57BL/6J (strain#:000664) mice from The Jackson Laboratory were used in all experiments. To reduce confounding variability, littermates were used whenever possible. All mice were housed in the same animal room under a 12-hr light-dark cycle with *ad libitum* access to food and water. Environmental conditions, including temperature and humidity, were kept consistent, with identical bedding and enrichment across all cages. Each cage housed 3-5 mice of the same sex to minimize social stress. Following tetrode implantation, animals were singly housed to ensure post-surgical recovery.

All procedures were conducted in compliance with the ARRIVE 2.0 guidelines to ensure methodological transparency and reproducibility. The experimental protocol was approved by the Institutional Animal Care and Use Committee (IACUC) at Nemours Children’s Health and was performed in accordance with the National Institutes of Health *Guide for the Care and Use of Laboratory Animals*.

### Handling and Habituation

All mice were handled daily by a single experienced experimenter for routine husbandry and any procedural injections. The same experimenter conducted all behavioral testing, reducing inter-experimenter variability that could influence animal responses. Mice were transferred to the testing room at least 60 minutes prior to the start of any behavioral task or recording to allow habituation. During testing, the experimenter remained outside the room to minimize potential stress or distraction.

### Early Life Seizure Mouse Model

Seizures begin on postnatal day 10 (p10). 20 total seizures were administered for 5 days from p10-p14. Wild type C57BL/6J mice were divided into three groups: control vehicle-treated, vehicle-treated ELS, and ACTH-treated ELS (N = 8 control vehicle-treated, N = 10 vehicle-treated ELS, N = 10 ACTH-treated ELS). An hour before the first seizure on each seizure day, mice received subcutaneous injections of their respective drug or vehicle. Animals took a one-hour break between each of their four daily seizure sessions. Animals were housed in individual sections of a custom-designed and built plexiglass chamber, which were linked to a central chamber that contained flurothyl. All sections of the chamber have equal access to the central chamber containing the flurothyl when the section doors are closed, but a sliding plastic panel cuts off flurothyl access once the section is opened for quick evacuation of the flurothyl.

### Flurothyl Induction

0.02mL of Flurothyl (Sigma-Aldrich) was dispensed into the central chamber on a filter paper and allowed to diffuse into the connected chambers containing individually placed animals. At incremental doses of 0.01mL or 0.02mL, flurothyl was given to elicit seizures with a minimum interval of one minute between each administration. The animal’s chamber was immediately evacuated upon the onset of a tonic-clonic seizure.

### Drug Administration

Mice in the ACTH group received subcutaneous injections with an ACTH in 5% gelatin solution at a dose of 150 IU/m^2^, diluted with 5% gelatin to a total volume of 0.1mL. Vehicle Control and Vehicle ELS littermates received 0.1mL subcutaneous injections of the same solution vehicle as their littermates. All mice received drug administration once daily, one hour prior to each day’s seizure inductions.

### Electrode Implantation

At p45, a custom 3D printed electrode with four drivable tetrodes was implanted through a 3mm burr hole into the prelimbic mPFC (+2.5 A/P and +0.5 M/L), a ground wire and a reference wire were placed above the cerebellar parenchyma, allowing us to record local field potentials (LFPs) and single units. The implant is glued and cemented to the skull followed by a 3 day recovery period. Electrodes were advanced 20μm for each recording session to ensure that different populations of neurons are being sampled during each recording session. After identifying and recording PFC cells during baseline, PFC cells are recorded during the performance of behavioral tasks.

### Single Neuron Recording

Electrode signals were preamplified and transmitted to the Neuralynx recording system (Neuralynx, Bozeman, MT). Single neuron action potentials are identified after the signal is filtered between 500-9000 Hz and thresholded for 3× root mean square (RMS) noise. Action potentials were then clustered and evaluated for quality control using L-ratio and isolation distance in SpikeSort3D. Number of recorded cells for each group during baseline is the following: control group (n=8, 149 cells), vehicle-treated ELS group (n=10, 133 cells) and ACTH-treated ELS group (n=10, 125 cells). The number of recorded cells for each group during fear extinction is the following: control group (n=7, 65 cells), vehicle-treated ELS group (n=7, 69 cells) and ACTH-treated ELS group (n=6, 59 cells).

### Generalized Linear Modeling

The clustered action potentials were used for further analysis through a generalized linear model (GLM). The GLM is used to quantify in vivo neural dynamics in healthy and early life seizure mice. Through the GLM, we are able to investigate the temporal modulation of neurons(*43*, *44*). Temporal modulation investigates how the neuron firing is modulated over time with respect to LFP firing. Post-spike filters (PSFs) estimate the effect of past spikes on current spike probability or fine spike timing and it is crucial for understanding the temporal dynamics of neuronal firing.

### Post-spike filters (PSFs) generation

A PSF is a mathematical function that represents the probability density of a neuron firing an action potential or spike as a function of time after it has previously fired(*43*, *44*). To generate PSFs, we started by analyzing the spike train of each neuron. Given that the refractory period of a neuron is approximately 2 ms, we used a 1 ms bin size to create a binned spike train, ensuring no more than one spike per bin. This binned spike train is modeled as a Poisson process, from which we derive the probability density function that maximizes the likelihood of observing the given spike train. To account for post-spike rate modulation (auto-correlation), we modeled the spike rate λ (t) as:

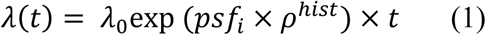

where (𝜌^ℎ𝑖𝑠𝑡^) denotes the spiking history of the neuron, and (psf_i_) is the post-spike filter encoding the firing rate modulation after a spike. The filters were parameterized using 10 raised cosine basis functions and include an immediate post-spike impulse to capture the refractory period(*42–44*). A ridge penalty was then added to the log-likelihood function to avoid overfitting(*82*). The resulting PSFs are computed for a maximum time bin of 662 ms, resulting in a 662-component vector for each post-spike filter. This process is implemented using Matlab.

After PSFs are implemented, principal component analysis is performed to establish the first principal component (PC1) to investigate fine spike timing heterogeneity within the recorded brain region; thus, providing a reliable readout of the neural network.

### Principal Component Analysis (PCA)

Principal Component Analysis (PCA) is a statistical technique used to reduce the dimensionality of large datasets while capturing as much variability as possible. After generating the post-spike filters (PSFs) for each neuron, PCA is applied to these filters to identify the main patterns of variance in the neuronal firing data. The PSFs, represented as 662-component vectors, form a high-dimensional dataset that is complex to analyze directly. PCA transforms this dataset into a new coordinate system defined by the principal components (PCs), which are orthogonal directions capturing the maximum variance in the data.

The first principal component (PC1) captures the largest amount of variance. By projecting the PSFs onto these principal components, we can reduce the dataset to a lower-dimensional space, typically focusing on the first few PCs that capture the most significant patterns. This reduction simplifies the analysis and visualization of the data, allowing us to investigate fine spike timing heterogeneity within the recorded brain region. The PCA-transformed data provides a reliable readout of the neural network’s activity, highlighting key differences and similarities in neuronal firing patterns across different conditions or groups. This process is implemented using Matlab, ensuring precise and efficient computation of the principal components.

### PSTH Construction and Firing Rate Percentage Change Analysis

To analyze neuronal firing rate dynamics during fear extinction learning (See section 15 below), we constructed peri-stimulus time histograms (PSTHs) and computed the percentage change in firing rate relative to baseline for two segments of the extinction session: the first 10 tones (early learning phase) and the last 10 tones (late learning phase). Baseline firing rates were calculated using spike timestamps within the pre-tone period preceding the onset of the first tone. The firing rate during this baseline period was determined by dividing the number of spikes by the duration. This baseline (pre-tone) firing rate served as the reference point for calculating percentage changes in subsequent analyses. To construct the PSTH, spike data were segmented into 3-second time bins corresponding to tone and no-tone periods.

For each time bin, the percentage change in firing rate relative to baseline was calculated using the formula:

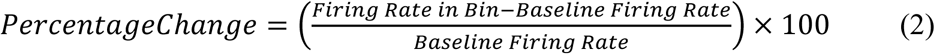

In addition to bin-wise PSTH analysis, we calculated the overall percentage change in firing rate for each tone and no-tone segment. PSTHs were visualized as bar graphs, with firing rate values plotted for each bin. The analysis was implemented in Python utilizing python NumPy, Matplotlib, and Pandas packages.

### Graph Metrics Analysis for Baseline and Fear Extinction

To assess neural activity and brain network evolution during the baseline and fear extinction sessions, we implemented a graph metrics-based approach. Graphs were constructed independently for the 600-second baseline session, as well as for the first and last 600-second segments of the fear extinction task. In each graph, nodes represented individual neurons, and weighted edges represented the functional connectivity between neuron pairs.

Firing rates were first computed using non-overlapping 10-second bins, resulting in 60 time points per 600-second segment. To ensure comparability between neurons with different firing rate scales and to minimize the dominance of high-firing neurons in the dot product computation, we applied min-max normalization to each neuron’s binned firing rate vector. This transformation scaled each neuron’s activity to a range of [0, 1] across time, preserving the temporal structure while standardizing the magnitude.

Functional connectivity between each neuron pair was quantified using the dot product of their min-max normalized firing rate vectors, calculated over a sliding window of three consecutive time bins, and advanced one bin at a time. The mean dot product across all sliding windows was used as the final edge weight. These weighted graphs were then used to compute various graph-theoretic metrics to capture the organization and evolution of network connectivity across conditions.

**Edge weights** represent the functional connectivity strength between pairs of neurons that were computed as:

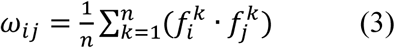

where f^k^_i_ and f^k^_j_ are the firing rates of neurons i and j in window k, and n is the total number of sliding windows. Higher edge weights indicate stronger functional connectivity between the neuron pairs, reflecting greater synchronization of their activity.

**Weighted degree centrality** reflects the overall connectivity of a neuron within the network, considering both the number and strength of its connections. For a neuron i, it is defined as:

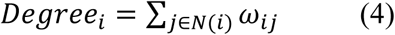

where w_ij_ is the weight of the edge between neurons i and j, and N(i) is the set of neighbors of neuron i.

To enable comparison across networks of different sizes, the weighted degree centrality was normalized:

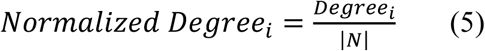

where ∣N∣ is the total number of neurons in the network.

Higher values of weighted degree centrality indicate neurons that are highly connected within the network, both in terms of the number and strength of connections, emphasizing their central role in network communication.

**Weighted clustering coefficient** quantifies the tendency of neurons to form densely interconnected clusters, was computed as:

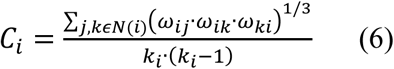

where w_ij_, w_jk_, and w_ki_ are the weights of the edges forming a triangle involving neuron i, k_i_ is the degree of neuron i (number of neighbors), and N(i) is the set of neighbors of neuron i.

Higher values of C_i_ indicate that the neuron participates in stronger and denser clusters within the network.

**Eigenvector centrality** was used to quantify the influence of individual neurons within the network, determined by their connectivity to other influential neurons. This metric assigns a score to each node (i) based on the centrality of its connected nodes, defined as:

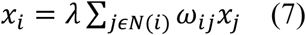

where x_i_ represents the eigenvector centrality of node i, w_ij_ is the weight of the edge between nodes i and j, N(i) is the set of neighbors of node i, and λ is the largest eigenvalue of the adjacency matrix. The equation is solved iteratively, normalizing the centrality values such that higher scores indicate nodes that are connected to other highly central nodes. Eigenvector centrality was calculated using the NetworkX Python library, which applies spectral decomposition methods to compute the centrality scores.

### Network Efficiency Metrics

#### Global Efficiency

Global efficiency (E_glob_), assessing the integration and efficient information transfer across the network, was computed as the average inverse shortest path length between all node pairs:

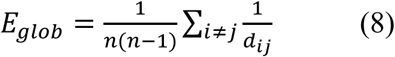

where d_ij_ is the shortest path length between nodes i and j. Nodes that are not connected (i.e., d_ij_=∞) were excluded from the calculation. Higher global efficiency values indicate a more integrated and efficient network.

#### Local Efficiency

Local efficiency (E_loc_) measures the fault tolerance and robustness of network communication within localized node neighborhoods:

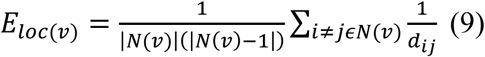

where N(v) is the set of neighbors of node v, and d_ij_ is the shortest path length between nodes i and j within the subgraph induced by N(v).

The overall local efficiency of the graph is the average of E_loc_(v) over all nodes:

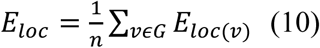

#### Analysis Workflow

Weighted, undirected graphs were constructed for each animal and segment using normalized edge weights. To account for the importance of stronger connections, weights were inverted (1-weight) to represent path lengths. Global and local efficiency were computed using the NetworkX library in Python, and all calculations were based on the largest connected component of each graph to ensure consistency.

### Graph Neural Network (GNN) Analysis for Predicting Behavioral Outcomes

We implemented a Graph Neural Network (GNN) using a Graph Attention Network (GAT) architecture to predict behavioral outcomes (percentage freezing) from neuronal network features and functional connectivity during the fear extinction task. Neuronal features included the firing rate, mean interspike interval (mISI), maximum ISI (max ISI), minimum ISI (min ISI), coefficient of variation (CV) of ISI, and principal component (PC1 and PC2) scores of the post-spike filters. Graphs were constructed for the first and last segments of the extinction session.

The GAT model consisted of three attention layers: two hidden layers using rectified linear unit (ReLU) activations and a final output layer with a sigmoid activation output scaled from 0 to 100% to match the behavioral outcome (percentage freezing). Model regularization techniques included dropout (optimized range: 0.1–0.5), weight decay (set at 1e-5), and a cosine annealing learning rate scheduler to enhance generalization and prevent overfitting. Hyperparameter optimization with hidden channel sizes (64, 128, 256), dropout rates, learning rates (0.01, 0.001, 0.0001), and attention heads (1–4) was performed using Optuna’s grid search sampler, evaluating different combinations of hyperparameters. For robust optimization, optimal hyperparameters were selected based on minimal combined validation loss and maximal R² scores through a 5-fold cross-validation scheme.

After optimization, model performance was rigorously assessed using Leave-One-Out Cross-Validation (LOOCV), ensuring robust generalization to unseen animals. Training efficacy and model convergence were monitored by plotting average training and validation loss curves across folds **(Supplementary Fig. 3)**. Predictive performance was assessed using the coefficient of determination (R²), computed as:

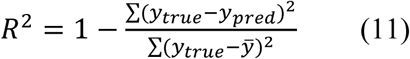

where 𝑦_𝑡𝑟𝑢𝑒_ denotes actual freezing percentage, and 𝑦̅ denotes their mean. An R² greater than 0 indicates that the model explains variability better than predicting the mean outcome. An R² of 0 indicates predictions are no better than the mean, while a negative R² indicates the model performs worse than predicting the mean.

To interpret model predictions, feature importance scores were derived using the GNNExplainer algorithm, which assigns feature importance scores based on their influence on predictions.

As a comparative baseline, we employed linear regression with LOOCV using average neuronal features per animal, as this model cannot incorporate node-level features directly. Additionally, we performed Generalized Estimating Equations (GEE) analyses to identify significant features associated with behavioral outcomes across neurons, facilitating comparison between methods.

### Open Field Task

At p50, mice were placed in a 46 cm x 46 cm squared arena in an isolated room and allowed to explore freely for 10 minutes. Sessions were video recorded while baseline single-unit recordings were performed.

### Fear Conditioning

At p55, mice (n=8 controls, n=10 vehicle-treated ELS and n=10 ACTH-treated ELS) performed a 2-day fear conditioning task.

Fear acquisition takes place on day 1. During fear acquisition, mice are individually placed in an operant box. During the acquisition task, mice experience four tone-shock pairings (28s tone followed by a 2s 0.25mA foot shock).

Fear extinction takes place on day 2. During fear extinction, mice are individually placed in the same operant box. During the extinction task, thirty 30s tones are presented without shocks at random intervals. Mice are recorded for PFC cells during fear extinction.

During both days, mice are video recorded and freezing time is quantified using ANY-maze. Scores were analyzed as a moving score ranging between 0 (total freezing) and 1 (total movement) or percentage freezing (0-100%).

### Statistical Analysis

All behavior, electrophysiology, and neural network metrics data were analyzed using a generalized estimating equation (GEE) model, a multivariable, repeated-measures, regression model. This approach allows for modeling data with multiple measures per animal and adjusts for potential confounding variables like sex and litter. Sex, genotype and group were used as factors. Sex and litter were not significant and were therefore removed from the final analyses. Non-significance is defined as p>0.05; p values for all significant results are reported in the results section.

To ensure that the study was adequately powered to detect group differences, a post hoc power analysis was conducted for each generalized estimating equation (GEE) model using G*Power(Faul et al., 2007). Effect sizes (Cohen’s f^2^) were calculated from the Wald Chi-Square (χ²) values reported in the GEE output using the formula f^2^=χ²/N, where N represents the total number of animals analyzed per model. For the dataset (N = 26), the Wald Chi-Square was χ² = 13.546 (df = 2, p < 0.001), yielding an effect size of f² = 0.521. This resulted in an 87.88% power at α = 0.05, indicating a high likelihood of detecting true group differences.

## Acknowledgments

Authors would like to acknowledge Pravin Wagley for their support.

## Funding

NIH NINDS 5K22NS104230 (AH) NIH NINDS 5R01NS134491 (AH)

R21NS117112/GF/NIH HHS/United States (RS)

## Author contributions

Designed experiments: MRK, JMM, RCS, and AEH

Conducted experiments: MRK, KAR, MO

Analyzed data: MRK, PJ, JMM and AEH

Writing: MRK, RCS, JMM and AEH

## Data and materials availability

Requests for further information and resources should be directed to and will be fulfilled by the lead contact, Mohamed R. Khalife.

All data reported in this paper will be shared by the lead contact upon request. All original code has been deposited at Zenodo at doi.org/10.5281/zenodo.15492107, doi.org/10.5281/zenodo.15492093, doi.org/10.5281/zenodo.15492126, doi.org/10.5281/zenodo.15492208, and is publicly available as of the date of publication.

Any additional information required to reanalyze the data reported in this paper is available from the lead contact upon request.

## Supplementary Materials

**Fig. S1.**
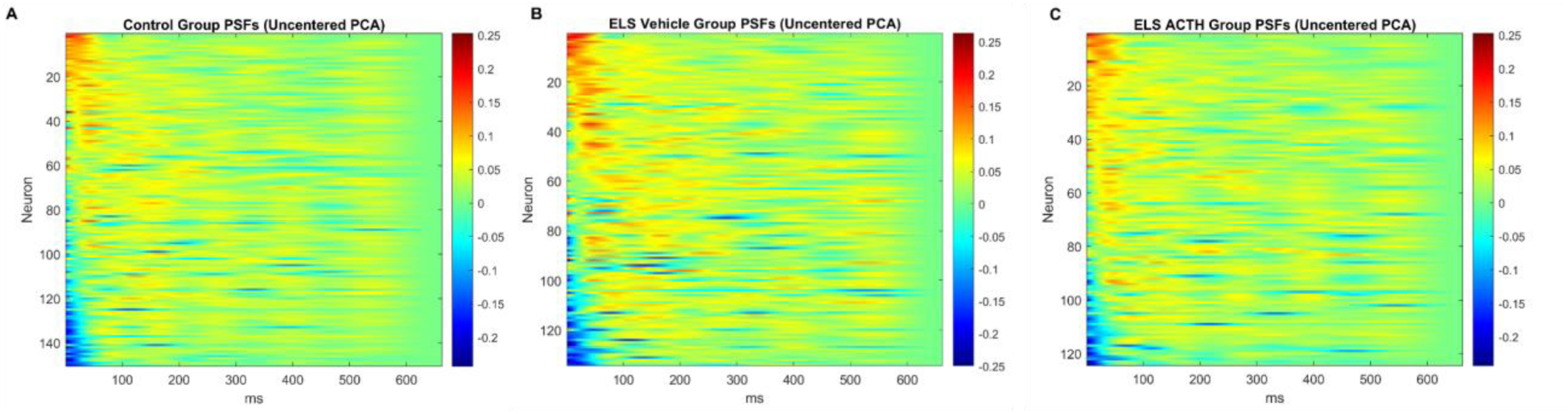
Heatmaps of post-spike filters (PSFs) sorted by PC1 score across groups. Post-spike filters (PSFs) from individual neurons are visualized as heatmaps and sorted by descending PC1 score within each group. Each row represents one neuron’s normalized PSF over a 662 ms window following a spike, capturing its temporal modulation of excitability. Control neurons show heterogeneous but smooth post-spike decay patterns with moderate early beta-range and weak theta-range modulation **(A)**. Vehicle-treated ELS neurons display prominent structure at around 42 and 120 ms, consistent with exaggerated beta and theta locking **(B)**. ACTH-treated ELS neurons show a more variable and less structured temporal profile, close to the control group **(C).** These heatmaps highlight group-level differences in temporal coding and support the main findings presented in Figure 3.

**Fig. S2.**
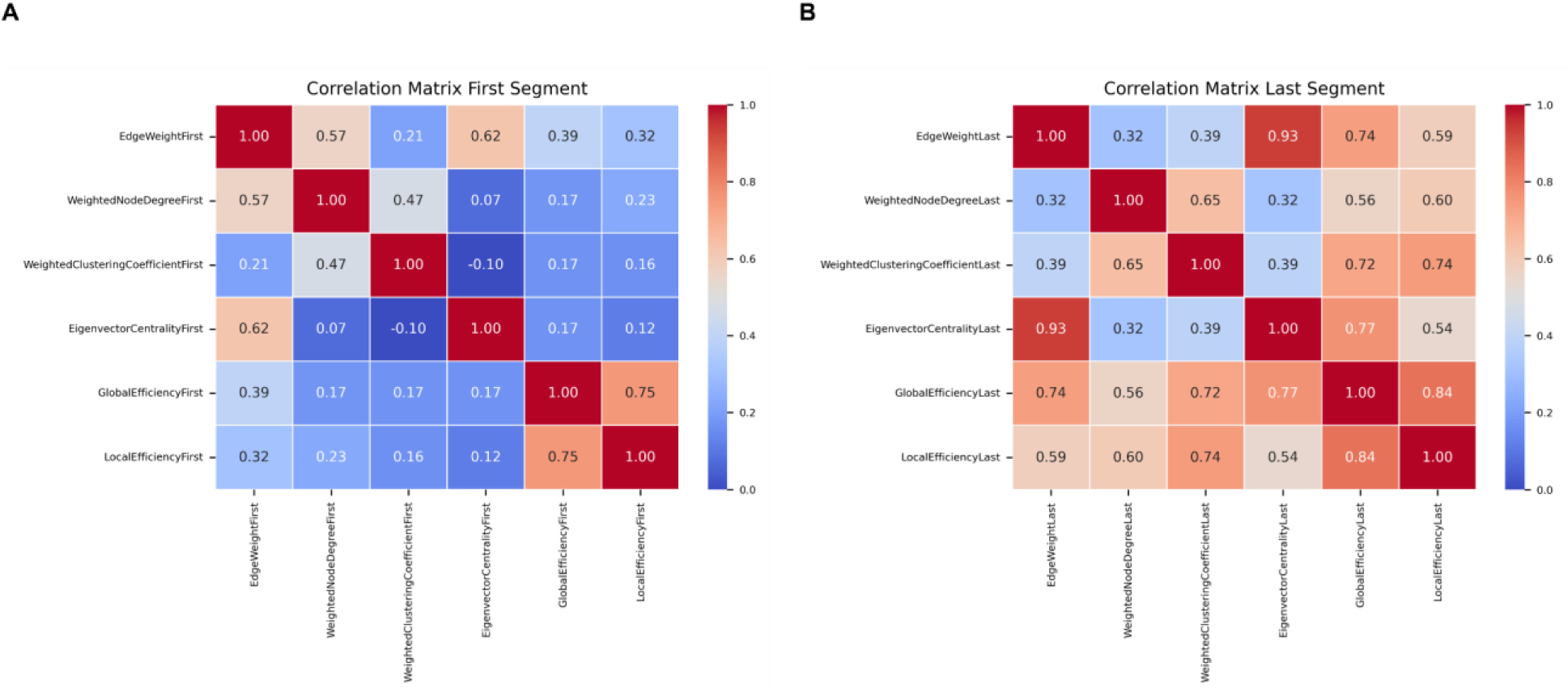
Spearman Correlation matrices of network metrics during fear extinction. Correlation matrices depicting associations between key network metrics at the first segment (left panel) and last segment (right panel) of fear extinction. Metrics include Edge Weight, Weighted Degree Centrality, Weighted Clustering Coefficient, Eigenvector Centrality, Global Efficiency, and Local Efficiency. During the initial segment, correlations were moderate, with edge weight correlating most strongly with weighted degree centrality. In the final segment, correlations among network metrics became notably stronger, highlighting the interdependent evolution of functional connectivity and network efficiency throughout extinction learning. Values indicate correlation coefficients, with color intensity proportional to correlation strength (red indicates positive correlations; blue indicates weak or negative correlations).

**Fig. S3.**
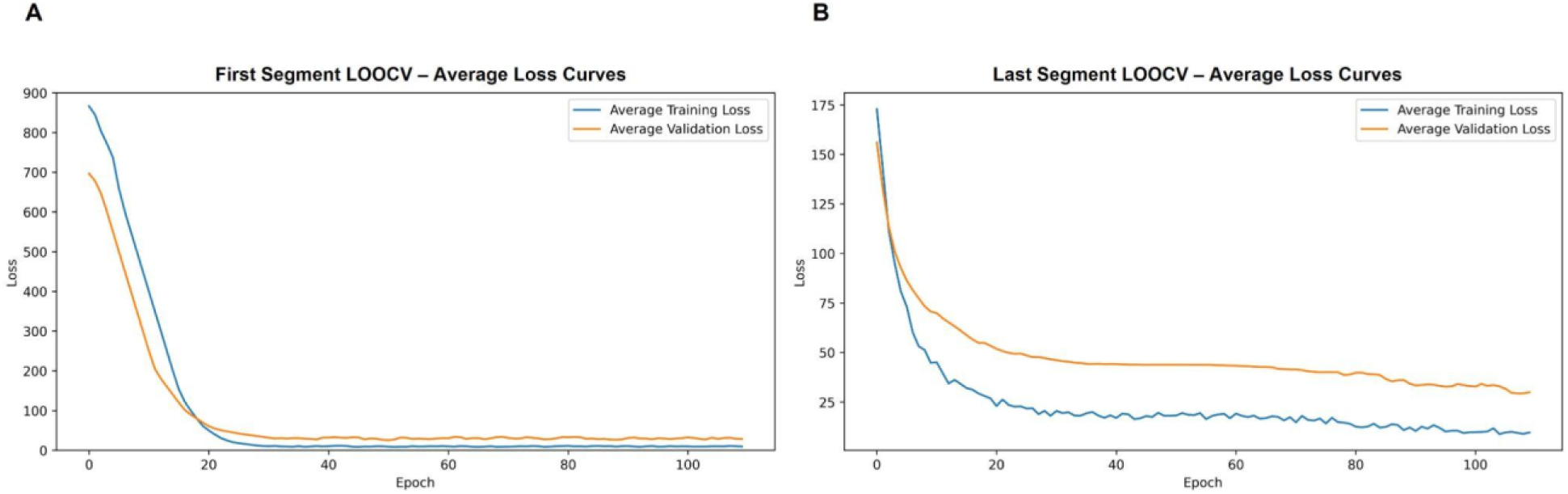
Average Training and Validation Loss Curves. Loss curves represent average training and validation losses across all Leave-One-Out Cross-Validation (LOOCV) folds for the GAT models trained to predict freezing behavior during the first segment (left panel) and last segment (right panel). The decrease and stabilization of both training and validation losses indicate successful optimization and model convergence.

**Table S1.**
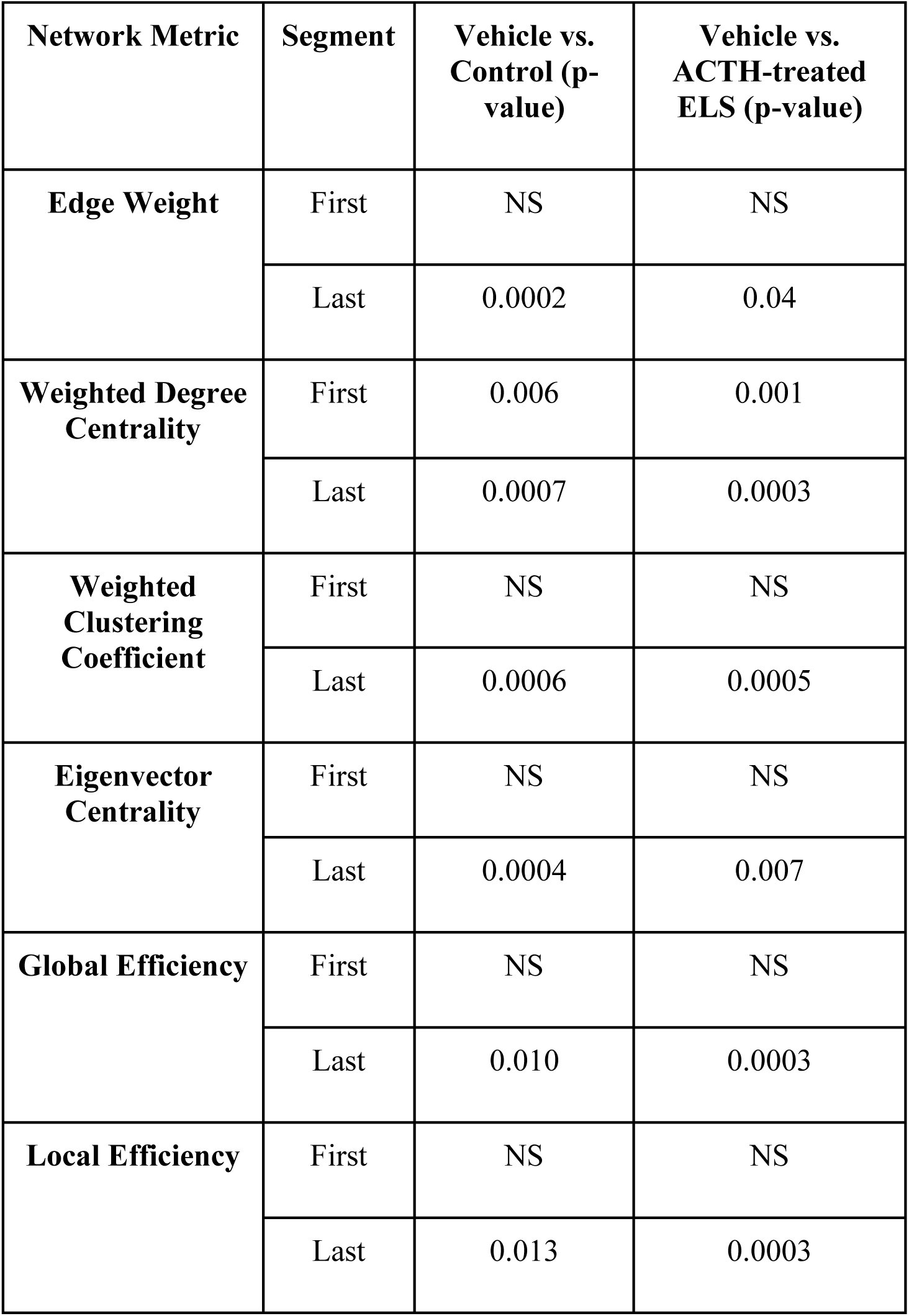
Summary of Network Metrics and Statistical Comparisons Across Groups. NS designates non-significance.

