## Supplemental Figures S1-S3 and Table 1 for "Normalization of Prefrontal Network Dynamics Prevents Cognitive Impairments After Developmental Insult"

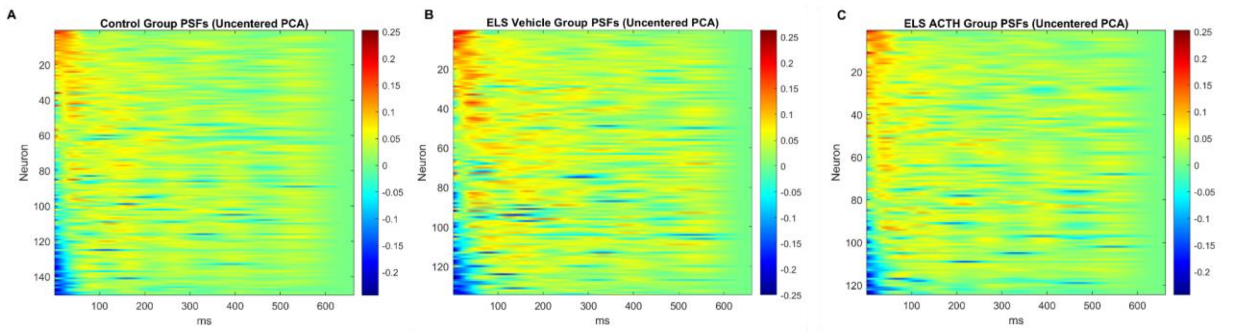


**Fig. S1.**

**Heatmaps of post-spike filters (PSFs) sorted by PC1 score across groups.** Post-spike filters (PSFs) from individual neurons are visualized as heatmaps and sorted by descending PC1 score within each group. Each row represents one neuron's normalized PSF over a 662 ms window following a spike, capturing its temporal modulation of excitability. Control neurons show heterogeneous but smooth post-spike decay patterns with moderate early beta-range and weak theta-range modulation **(A)**. Vehicle-treated ELS neurons display prominent structure at around 42 and 120 ms, consistent with exaggerated beta and theta locking **(B)**. ACTH-treated ELS neurons show a more variable and less structured temporal profile, close to the control group **(C).** These heatmaps highlight group-level differences in temporal coding and support the main findings presented in Figure 3.

.

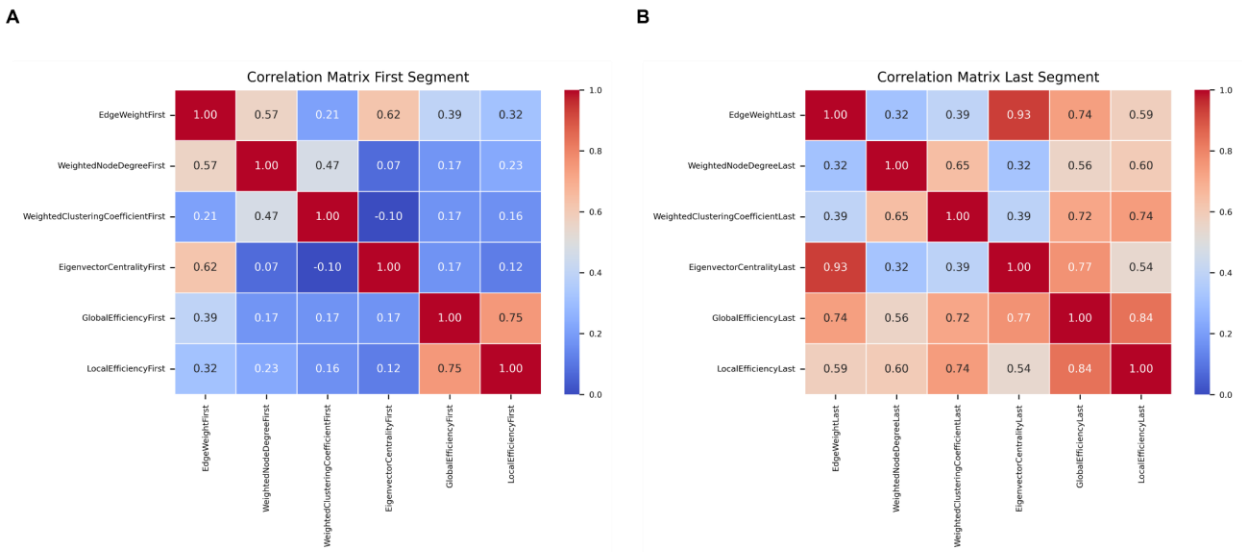


**Fig. S2.**

**Spearman Correlation matrices of network metrics during fear extinction.** Correlation matrices depicting associations between key network metrics at the first segment (left panel) and last segment (right panel) of fear extinction. Metrics include Edge Weight, Weighted Degree Centrality, Weighted Clustering Coefficient, Eigenvector Centrality, Global Efficiency, and Local Efficiency. During the initial segment, correlations were moderate, with edge weight correlating most strongly with weighted degree centrality. In the final segment, correlations among network metrics became notably stronger, highlighting the interdependent evolution of functional connectivity and network efficiency throughout extinction learning. Values indicate correlation coefficients, with color intensity proportional to correlation strength (red indicates positive correlations; blue indicates weak or negative correlations).

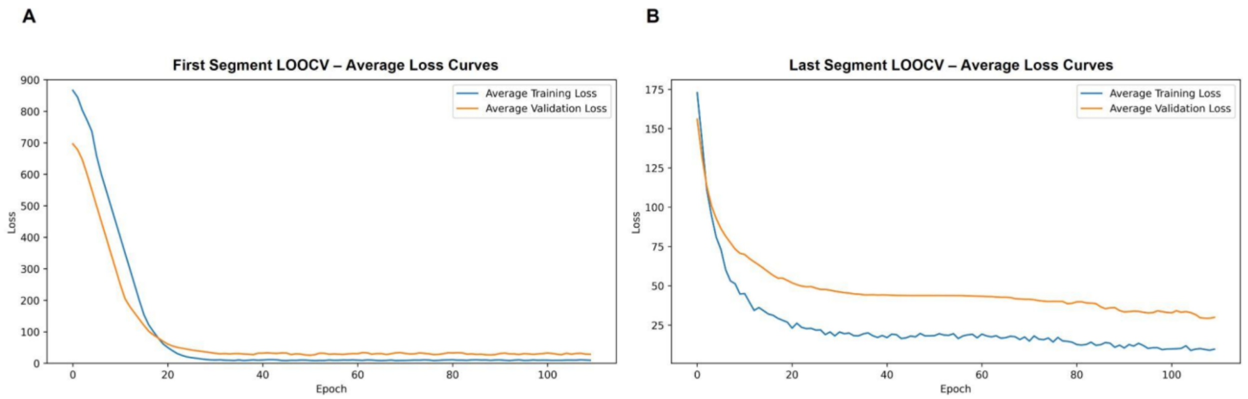


**Fig. S3.**

**Average Training and Validation Loss Curves.** Loss curves represent average training and validation losses across all Leave-One-Out Cross-Validation (LOOCV) folds for the GAT models trained to predict freezing behavior during the first segment (left panel) and last segment (right panel). The decrease and stabilization of both training and validation losses indicate successful optimization and model convergence.

| Network Metric | Segment | Vehicle vs. Control (p-value) | Vehicle vs. ACTH-treated ELS (p-value) |
|---|---|---|---|
| **Edge Weight** | First | NS | NS |
| | Last | 0.0002 | 0.04 |
| **Weighted Degree Centrality** | First | 0.006 | 0.001 |
| | Last | 0.0007 | 0.0003 |
| **Weighted Clustering Coefficient** | First | NS | NS |
| | Last | 0.0006 | 0.0005 |
| **Eigenvector Centrality** | First | NS | NS |
| | Last | 0.0004 | 0.007 |
| **Global Efficiency** | First | NS | NS |
| | Last | 0.010 | 0.0003 |
| **Local Efficiency** | First | NS | NS |
| | Last | 0.013 | 0.0003 |

**Table S1.**

**Summary of Network Metrics and Statistical Comparisons Across Groups. NS designates non-significance**
